# Sex-specific differences in rotarod performance and type 1 cannabinoid receptor levels in a rat model of traumatic brain injury treated with Δ^9^-tetrahydrocannabinol

**DOI:** 10.1101/2022.04.11.487790

**Authors:** Tallan Black, Ayat Zagoog, Andrew J Roebuck, Quentin Greba, J. Patrick Neary, John G. Howland, Robert B. Laprairie

## Abstract

Traumatic brain injuries (TBI) remain one of the leading causes of death and disability world-wide. One emerging area of TBI research is the involvement of the endocannabinoid system (ECS) in response to TBI. Endogenous cannabinoids modulate inflammation, pain, anxiety, and neurotransmitter release through the activation of the cannabinoid receptors CB1R and CB2R. CB1R and CB2R are activated by exogenous cannabinoids such as Δ^9^-tetrahydrocannabinol (THC) found in *Cannabis sativa*. As public perceptions change in the wake of *Cannabis* legalization, research into the potential harmful and therapeutic effects of THC following TBI deserve exploration. In this preliminary study, we investigated sex differences in behavioral effects, CB1R abundance, and cytokine profiles in a rat model of moderate TBI treated with 1 mg·kg^-1^ THC (*i*.*p*.). Neither TBI nor THC treatment altered catalepsy, body temperature, nociception, or spontaneous alternation as measured in the y-maze. TBI reduced male rotarod performance in both vehicle and THC-treated groups, and THC treatment decreased performance in Sham-TBI rats when compared to vehicle controls. Female rats that received a TBI and THC exhibited lower relative CB1R density when compared to the Sham-TBI+THC group. TBI was associated with reduced interleukin-4 in males; THC increased interleukin-6 in TBI males compared to Sham-TBI. These preliminary results highlight fundamental sex differences in the response of the ECS following TBI. Our results indicate the need for further investigation of the ECS and phytocannabinoids post-TBI in both acute and chronic phases.

**Significance Statement:** The endogenous cannabinoid system is a potential target in the pathophysiology and treatment of traumatic brain injury (TBI). In this study we observed TBI reduced rotarod performance in male rats only and performance was not affected by THC. Female rats the received THC and TBI displayed lower cortical cannabinoid receptor 1 levels. These early results showcase sex differences in rodent models of TBI and the endogenous cannabinoid system.

## Introduction

Traumatic brain injury (TBI) is a complex pathophysiological condition with overlapping and ongoing symptomology. TBIs most commonly result from falls, sports-related injuries, car accidents, or physical assaults, and are usually accompanied by loss of consciousness, post-traumatic amnesia, as well as persistent cognitive and neurological deficits (Menon et al., 2010; Bramlett and Dietrich, 2015). Although pharmacological interventions ranging from lithium for neuroprotective and mood stabilizing effects to erythropoietin for anti-inflammatory and angiogenic effects have been explored for TBI with varying success, one emerging area of TBI research is the involvement of the endocannabinoid system (ECS) in response to brain injury (Ray et al., 2002; Diaz-Arrastia et al., 2014; Singh and Neary, 2020).

TBI is characterized by numerous pathophysiological changes within the brain that are triggered by a primary structural injury (Ray et al., 2002; Barkhoudarian et al., 2011; Xiong et al., 2014; DeKosky and Asken, 2017; Pearn et al., 2017; Sulhan et al., 2020). The secondary injury cascade involves widespread cellular, molecular, and biochemical changes that develop following the primary injury (Giva and Hovda et al., 2014; Currie et al., 2016; Jassam et al., 2017; Simon et al., 2017; Thibeault et al., 2019; Moore et al., 2020). The secondary injury evolves from minutes to years post-injury due to neuronal hyperexcitability, glial cell dysregulation, lipid degradation, cerebral edema, nitric oxide synthesis, and widespread neuroinflammation, which in turn leads to tissue damage, cellular atrophy, and the accumulation and aggregation of harmful biomolecules such as the amyloid precursor and tau protein (McDonald et al., 2002; Farooqui, 2010; Morganti-Kossmann et al., 2010; Kinoshita, 2016; DesKosky and Asken, 2017). Symptoms of short-term behavioral and cognitive impairments, and TBI-induced neurodegeneration, typically follow many of these brain regional-specific neurobiological alterations.

The type 1 and 2 cannabinoid receptors (CB1R, CB2R) are the predominant receptors of the ECS (Pertwee, 2010; Howlett and Abood, 2017). CB1R activation is considered neuroprotective because it depresses glutamatergic excitotoxicity (Marsicano et al., 2003), which in turn slows the production of reactive oxygen and nitrogen species and associated cell death (Schurman and Lichtman, 2017). CB2R activation within the CNS and periphery inhibits the release of pro-inflammatory cytokines, and this receptor’s activation is therefore associated with anti-inflammatory activity (Turcotte et al., 2016). THC, the main psychoactive component of *Cannabis*, is a partial agonist of CB1R and CB2R (McDonald et al., 2002). Human data are limited, but a 3-year retrospective study by Nguyen et al. (2014) of TBI patients who screened positively on urine toxicology for THC had a decreased mortality in comparison to patients who screened negatively. Preclinical data indicate that repeated administration of 1.25 mg·kg^-1^ (*i*.*p*.) THC after – but not before – repeated mild TBI in rats is associated with partial improvement in anxiety, and depression, and working memory deficits (Bhatt et al., 2020) and the treatment of male mice subjected to TBI with the CB2R agonist O-1966 decreases blood brain barrier disruption, decreases neuronal damage, and improves rotarod performance when compared to vehicle control (Amenta et al., 2012).

Based on the limited existing evidence connecting the ECS, TBI, and THC, the purpose of this preliminary study was to assess whether treatment of rats subjected to a moderate TBI (mTBI) with THC post-injury would affect behavioral and molecular deficits relative to non-injured rats.

## Materials and Methods

### Materials

All materials were from Sigma-Aldrich (Mississauga, ON), unless otherwise noted. Purified THC (98.9%) was purchased from Aurora Cannabis (Edmonton, AB, Canada).

### Animals

Thirty-six Sprague-Dawley rats of both sexes (8-12 weeks old) were used in this study (Charles Rivers Laboratories, Senneville, QC). Animals were acclimatized for a 7-day period without handling. The rats were housed in same-sex pairs with appropriate environmental enrichment, bedding materials, and *ad libitum* access to food and water. Animals were maintained on a standard 12:12 light dark cycle. Following initial acclimatization, animals were handled for 7 days prior to experimentation. All procedures and protocols described below were performed with approval from the University Animal Care Committee and Scientific Merit Review Committee for Animal-Based Research and are in keeping with the guidelines of the Canadian Council on Animal Care and the ARRIVE guidelines.

Three cohorts of 12 rats were randomly assigned into 4 experimental groups: (i) Vehicle [ethanol:emulphor:saline (1:1:18), *i*.*p*.] + Sham-TBI; (ii) Vehicle + TBI; (iii) THC (1 mg·kg^-1^) + Sham-TBI; (iv) THC + TBI. TBI was conducted as described below. Animals were anesthetized under isofluorane for 10±2 min in both Sham and TBI groups. A dose of 1 mg·kg^-1^ THC *i*.*p*. was administered 1 h post-TBI and was chosen as this dose approximates the ED_50_ for THC drug discrimination in the rat and other rodents (Zani et al., 2007; Järbe et al., 2014). On day 7, following the completion of *in vivo* data collection, all rats were euthanized by deep anesthesia using isoflurane (5%) for approximately 10 min – or until a pedal reflex could not be evoked – followed by decapitation. Whole brains were collected and stored at -80°C.

### TBI protocol

The mTBI model used was adapted from Mychasiuk et al. (2016) and Qin et al. (2018); this model is described as a ‘mixed model injury’. The mixed model injury is a variation of the weight drop and controlled cortical impact (CCI) models wherein animals are placed on foam or allowed to rotate or free fall following the weight drop or CCI (Mychasiuk et al., 2014). Helmet-like devices have been used to decrease the injury severity and modify the ratio of focal to diffuse injury type (Ma et al., 2019).

Following consultation with the University of Saskatchewan’s animal care committee and colleagues familiar with this mTBI model (Zani et al., 2007; Mychasiuk et al., 2014, 2016, Qin et al., 2018), the weight drop model was chosen due to its ability to produce, at low cost, a reliable and reproducible injury, producing similar behavioral deficits characterized by other more complex pre-clinical models (Amenta et al., 2012; Bhatt et al., 2020). Prior to TBI, rats were habituated to the procedure room for 15 min. To elicit a TBI, rats were individually anesthetized with 1.5 L·min^-1^ oxygen and isoflurane (3%) until their pedal reflex was no longer responsive. Following anesthesia, rats were then briefly transferred to a head cone and heating pad where fur along the dorsal cranium was shaved, and a stainless-steel disc (“helmet”) (10 mm × 2 mm) was fixed with dental cement on the sagittal mid-line between the inter-aural line and bregma to indicate the impact site. Animal body temperature was monitored by rectal thermometer throughout this procedure. Once the helmet was securely fastened, the rat was transferred to aluminum foil 20 cm above a foam pad, and placed in the prone position with their helmet positioned below the guide tube (Fig. 1). At this point, the animals ceased to receive anesthetic. The use of aluminum foil to suspend the rat allows for it to rotate 180° following impact of the weight being dropped, producing a more functional mechanism of injury (Mychasiuk et al., 2014). In total, anesthetic duration was 10±2 min for all animals. A 300 g weight was dropped from a height of 1 m impacting the “helmet” at the sagittal midline to produce a TBI (Fig. 1a). This weight corresponds to a force of approximately 2,900 N and is greater than weights used in previous studies (150 g and 200 g) but within the literature range for a rodent model of mTBI (Amenta et al., 2012; Mychasiuk et al., 2016; Ma et al., 2019; Bhatt et al., 2020). Of note, this force approximates the concussive forces applied to a human mandible of ∼2,100 N (Viano et al., 2012), Following impact, the rat was transported to a recovery cage placed on top of a heating pad with a pulse oximeter to monitor heart rate and blood oxygen content, and a rectal thermometer, until it regained consciousness (Extended Data Fig. 1-1a,b). No differences in animal weight were observed between groups 7 days after TBI or THC treatment (Extended Data Fig. 1-1c). Computed tomography scans showed no cranial fractures (Extended Data Fig. 1d). No differences in recovery time were observed across subjects and no mortalities occurred during this study (data not shown). A complete timeline of experimental TBI procedures, behavioural, and physiological measurements is provided in figure 1b.

**Figure 1.**
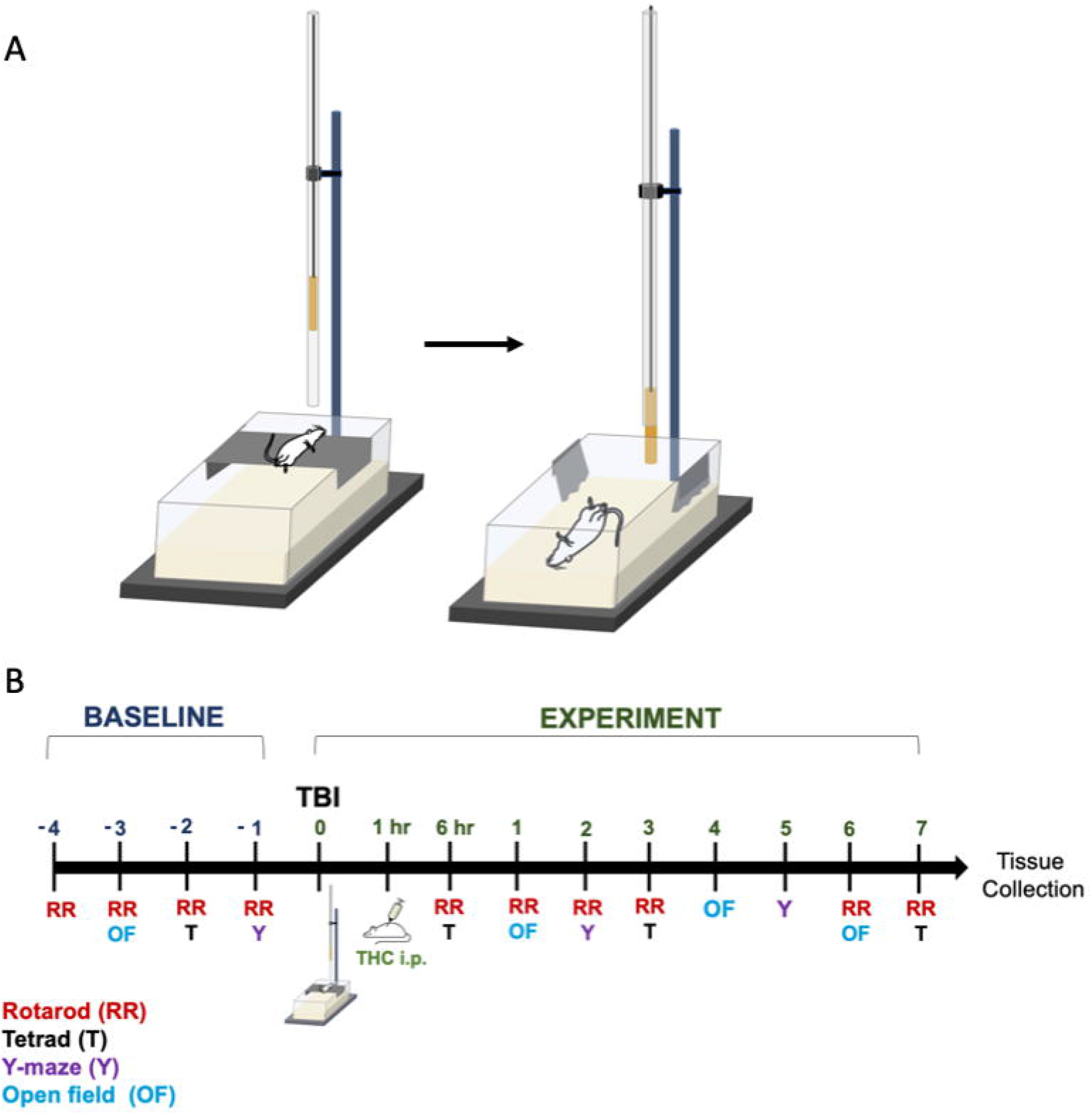
Modified Weight Drop Model. **(A)** Graphical representation of the TBI model used to emulate a combination of diffuse and focal injuries characteristics commonly seen in sports related and motor vehicle accidents. Figure created by author in reference to model adapted from (30). **(B)** Timeline representation of measurements and interventions performed in this experiment.

### Tetrad test

Tetrad tests were conducted to detect any acute or lasting cannabimimetic effects for treatment once with 1 mg·kg^-1^ THC, and are described below (Grim et al., 2016; Garai et al., 2020), 1 day before treatment (i.e., baseline), at 1 h, 3 days and 7 days post-TBI. To test catalepsy, rats were placed such that their forepaws rested on a 0.7 cm diameter bar 4.5 cm above a clean table. The duration spent holding the bar was recorded up to a 60 sec maximum. Time recording ceased when the rat either moved their forepaws off the bar or when the rat moved their head to the left or the right. Three trials were performed on each measurement day, and the mean time spent positioned on the bar correctly was used for analysis. Body temperature was measured with a rectal probe thermometer 10 min after catalepsy was tested. Anti-nociception was measured 15 min after catalepsy using the Tail Flick Analgesia Meter (Series 8, IITC Life Science, Woodland Hills, CA, USA). Rats were wrapped in a towel and positioned such that the heat light shone at 5 cm from the base of the tail. Time to tail flick was recorded up to a 20 sec maximum. The open field test (OFT) is a measure of locomotor activity and used to interpret levels of anxiety by making use of rodents’ aversion to open environments (Seibenhener and Wootten, 2015). Rats were placed in the center of the OFT apparatus (outer diameter: 1.5 m, inner diameter: 1 m). Locomotion was tracked using Ethovision XT (Noldus) for 10 min. All equipment for tests and others was thoroughly cleaned before and after use with 70% ethanol.

### Y-maze

Y-maze is a behavioral modality frequently used to assess the acute and chronic changes in spatial memory for both TBI and pharmacological investigation in rodents (Hughes, 2004). Animals were habituated to the treatment room for 1 day prior to data collection. Female rats were trained and tested before male rats each day. A baseline measure was established on day -1 pre-TBI. Rats were put into a different arm for each recording session as per the established timeline and recorded for 8 min through an overhead video camera. Videos were hand-scored and the percentage of spontaneous alternations as well as the number of entries into each arm were calculated. Observers were blinded to treatments. The percentage of alternation is equal to the number of correct alternations, divided by the total number of entries multiplied by 100. Videos were also analyzed for total locomotion using Ethovision XT. The entry arm was not counted in the total number of entries (Mandillo et al., 2008). One vehicle + Sham-TBI female was removed from the analysis due to violating test parameters by climbing out of the box.

### Rotarod

The rotarod is a test used to quantify vestibulo-motor function in rats following TBI by testing grip strength, coordination, and balance of the animal (Hamm et al., 1994). Rats were pre-trained on the rotarod (ROTO-ROD, Series 8 IITC Life Science) for 4 days prior to TBI (day -4 to -1) and tested on days 0, 1, 2, 3, 6 and 7 following TBI. Animals were habituated to the treatment room and instrument for 3 days prior to data collection. Rats were held in a neighboring room for a minimum of 10 min, and then habituated to the rotarod room for 5 min pre-test. Rotation was turned on with a smooth acceleration (2 rpm/5 sec) beginning from 4 rpm to a maximum of 36 rpm (achieved in 1 min 25 sec and maintained for the duration of the trial). The latency to fall was measured in sec to a maximum trial time of 5 min. Each animal was tested 3 times with a 5 min rest between trials. The mean of all 3 trials was used as the final score for each rat. The rotarod was thoroughly cleaned with the general virucide disinfectant PerCept^TM/MC^RTU.

### SDS-PAGE and western blot

Tissues were collected from animals euthanized on day 7 of the experiment (Fig. 1b). The TBI site was below the “helmet” along the sagittal midline. A section of cortex taken from below the injury site (or “helmet” for Sham animals) weighing approximately 100 µg (selected randomly from either the right or left hemisphere) was dissected from euthanized animals and homogenized (Rotor-Stator, OMNI International) to either be used for SDS-PAGE or cytokine analysis. The “helmet” was placed along the midline and therefore neither side is considered ipsi- or contralateral. Protein concentrations were quantified using the Pierce™ BCA protein assay kit (Thermofisher, Burlington, ON). Seventy µg of tissue homogenate was resolved on 10-20% Novex Tris-Glycine Mini Protein Gel (Thermo Fisher) for 20 min at 75 V, and 125V for 60 min. Protein was then transferred on ice at 25 V for 120 min onto 0.45 µm nitrocellulose membranes. Membranes were blocked with 20% Odyssey Blocking Buffer (Li-Cor, Lincoln, NE) for 1 h before being incubated overnight at 4°C with primary antibodies directed against the N-terminal extracellular region of the CB1R (1:500, rabbit polyclonal antibody, Cayman Chemical Company, Cat# 101500, Lot# 0580081-1) and βactin (1:10,000, mouse monoclonal antibody, Thermo Fisher, Cat# MA5-15739, Lot# UH285337) diluted in 20% Odyssey Blocking Buffer. Membranes were rinsed 6 times in 1X TBST before being incubated for 1 h at room temperature protected from light with Li-COR IR Dye^680^ (1:500) donkey anti-rabbit (Catalogue # 926-68073, Lot# D00303-13) and Li-COR IR Dye^800^ (1:500) β- actin donkey anti-mouse (Catalogue # 926-32212, Lot# D00226-15) secondary antibodies. Membranes were rinsed again 6 times in 1X TBST protected from light. Membranes were scanned using Odyssey Li-Cor (iS Image Studio 5.X). ImageJ (1.51) was used to perform densitometry analyses and quantification. Data are presented as relative ratios of CB1R to βactin levels. One female TBI + vehicle sample was excluded from analysis due to a lack of detectable protein. The CB1R antibody utilized here has previously been demonstrated to be specific to CB1R in western blotting with this membrane using CB1R blocking peptide and samples lacking CB1R (Roebuck et al., 2021).

### Cytokine analysis

Protein samples from cortex prepared as described in section 2.7 above were stored at -80°C. Frozen samples were sent to Eve Technologies and analyzed using the Rat Cytokine/Chemokine Array 27-Plex (BioPlex 200 Cytokine Array Assay Kit Source, Millipore Milliplex; Calgary, AB, Canada).

### Statistical analysis

Where sex-dependent differences appeared to be present, behavioral data were initially analyzed according to a four-way repeated measure (RM) analysis of variance test (ANOVA) using a 2×2×2×3, 4 or 7 (sex x injury x drug x time) randomized design. Data were separated by sex and analyzed according to a three-way 2×2×3, 4 or 7 (injury x drug x time) RM ANOVA for tests without sex-dependent differences. Where time was not significant, data was aggregated by time and sex and analyzed for injury and drug via two-way ANOVA 2×2 (injury x drug) randomized design. Molecular data were analyzed according to a three-way ANOVA using a 2×2×2 (sex x injury x drug) randomized design. Data for OFT (total movement and time in center) and rotarod were normalized as a measure of fold change from baseline for each subject. Normalization was performed to account for variability in performance between animals during baseline initial testing and account for scale differences. Sphericity was confirmed according to Mauchley’s test of sphericity. If data were not spherical, Greenhouse-Geisser-corrected values were used. Mauchley’s Correction for multiple comparisons was done using Tukey’s post-hoc analysis. Data is presented as mean ± standard error of the mean (SEM) from each group, *p < was considered statistically significant. All data was analyzed using GraphPad Prism 8.0 and IBM SPSS. All statistics are fully described and reported in Extended Data Tables 2-1 – 7-1.

## Results

### Tetrad

Assessments of catalepsy, body temperature, nociception, and movement in the OFT occurred on days -1 (baseline), 0 (day of TBI), 3, and 7. In the ring-holding test for catalepsy, no main effects were detected for injury, drug, time, or interactions therein for either males or females (Fig. 2A,B; Extended Data Table 2-1). With respect to body temperature, no main effects were detected for injury, drug, or interactions therein for either males or females (Fig. 2C,D; Extended Data Table 2-1). A main effect of time was observed for body temperature in males (p = 0.037) and females (p = 0.010), but Tukey’s post-hoc analyses did not reveal significance between time points (Fig. 2C,D; Extended Data Table 2-1). In the tail flick test for nociception, no main effects were detected for injury, drug, time, or interactions therein for either males or females (Fig. 2E,F; Extended Data Table 2-1).

**Figure 2.**
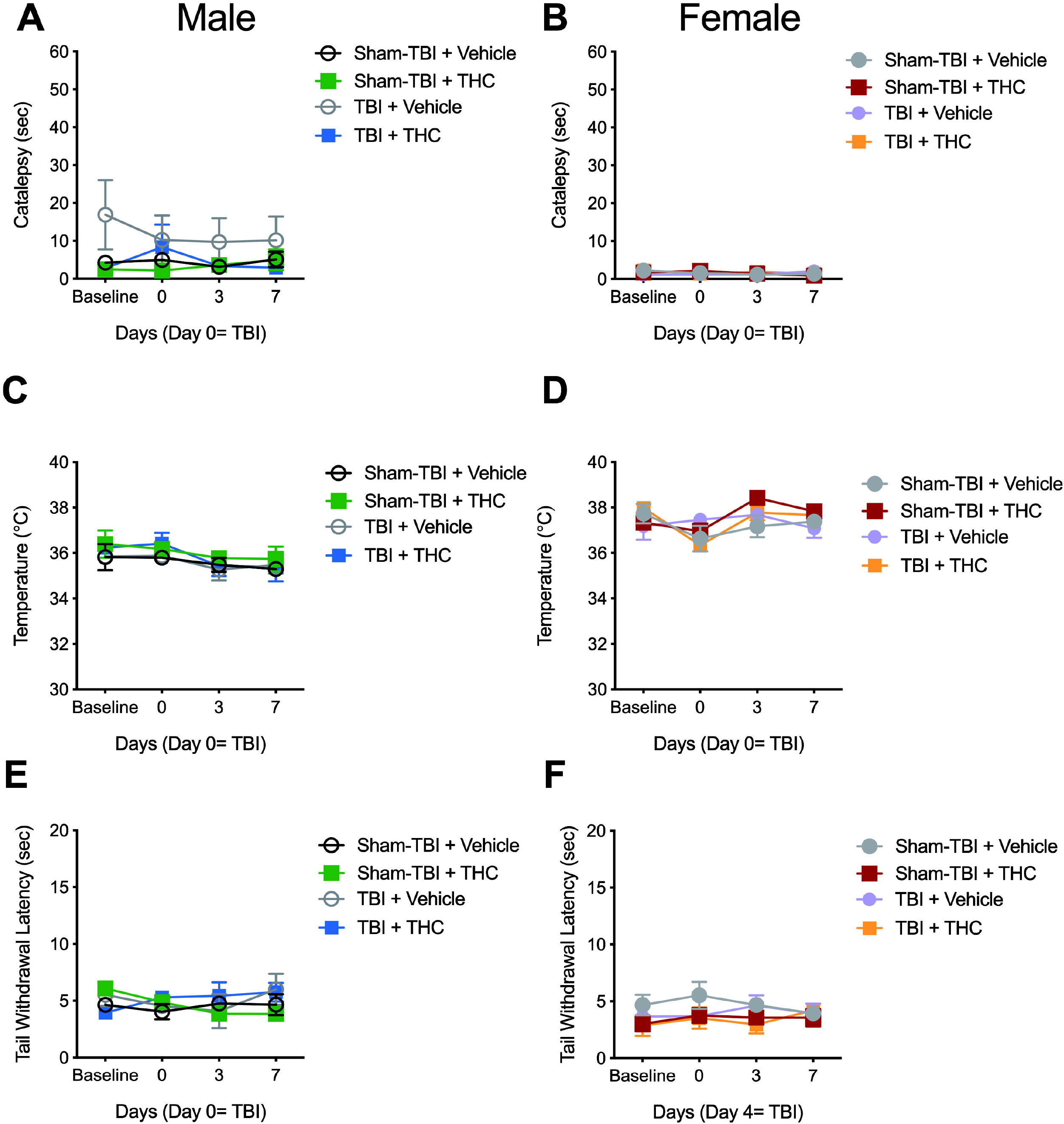
Assessment of catalepsy, body temperature, and nociception in rats following TBI and 1 mg·kg^-1^ THC treatment. Eighteen Sprague-Dawley rats of both sexes were administered a Sham-TBI or TBI and injected once with 1 mg·kg^-1^ THC *i*.*p*. or vehicle 1 h post-TBI. 1 mg·kg^-1^ THC *i*.*p*. nor TBI produced catalepsy in males **(A)** or females **(B)**. 1 mg·kg^-1^ THC *i*.*p*. nor TBI effected body temperature in males **(C)** or females **(D)**. 1 mg·kg^-1^ THC *i*.*p*. nor TBI effected nociception in males **(E)** or females **(F)**. Data presented as mean ± S.E.M. n=4-5 per group per sex. Statistical analyses were three-way RM ANOVA and described in Extended Data Table 2-1. Y-axes are set according to the minimum and maximum observable effects in each test (e.g. 60 sec for catalepsy).

In the OFT, total distance data were normalized to individual baseline scores in order to account for initial differences in total distance travelled by individual animals (Fig. 3; Extended Data Fig. 3-1). When either raw or normalized data were analyzed, no changes in total distance travelled were detected for sex, injury, time, or interactions (Fig. 3A,B). A main effect of drug was observed for total distance travelled, with THC-treated animals moving less than vehicle-treated animals (p = 0.024), but Tukey’s post-hoc analyses did not reveal significance between groups within days (Fig. 3A,B). Similar to distance travelled, time in the centre was normalized to individual baseline scores (Extended Data Fig. 3-1). No changes in time in the centre of the OFT were detected for injury or interactions (Fig. 3C,D). A main effect of sex was observed for time in the centre, with males spending more time in the centre than females (p = 0.040), but Tukey’s post-hoc analyses did not reveal significance between sexes within treatment groups (Fig. 3C,D). Similarly, main effects were observed for drug, with THC treatment associated with less time in centre than vehicle (p = 0.016) and time such that animals spent increasing time in the centre over the course of the experiment (p = 0.034), but Tukey’s post-hoc analyses did not reveal significance between treatment groups or time points, respectively (Fig. 3C,D). Therefore, neither TBI or THC changed performance in the tetrad, although males spent an increasing amount of time in the centre of the OFT regardless of treatment and likely due to repeated task exposure.

**Figure 3.**
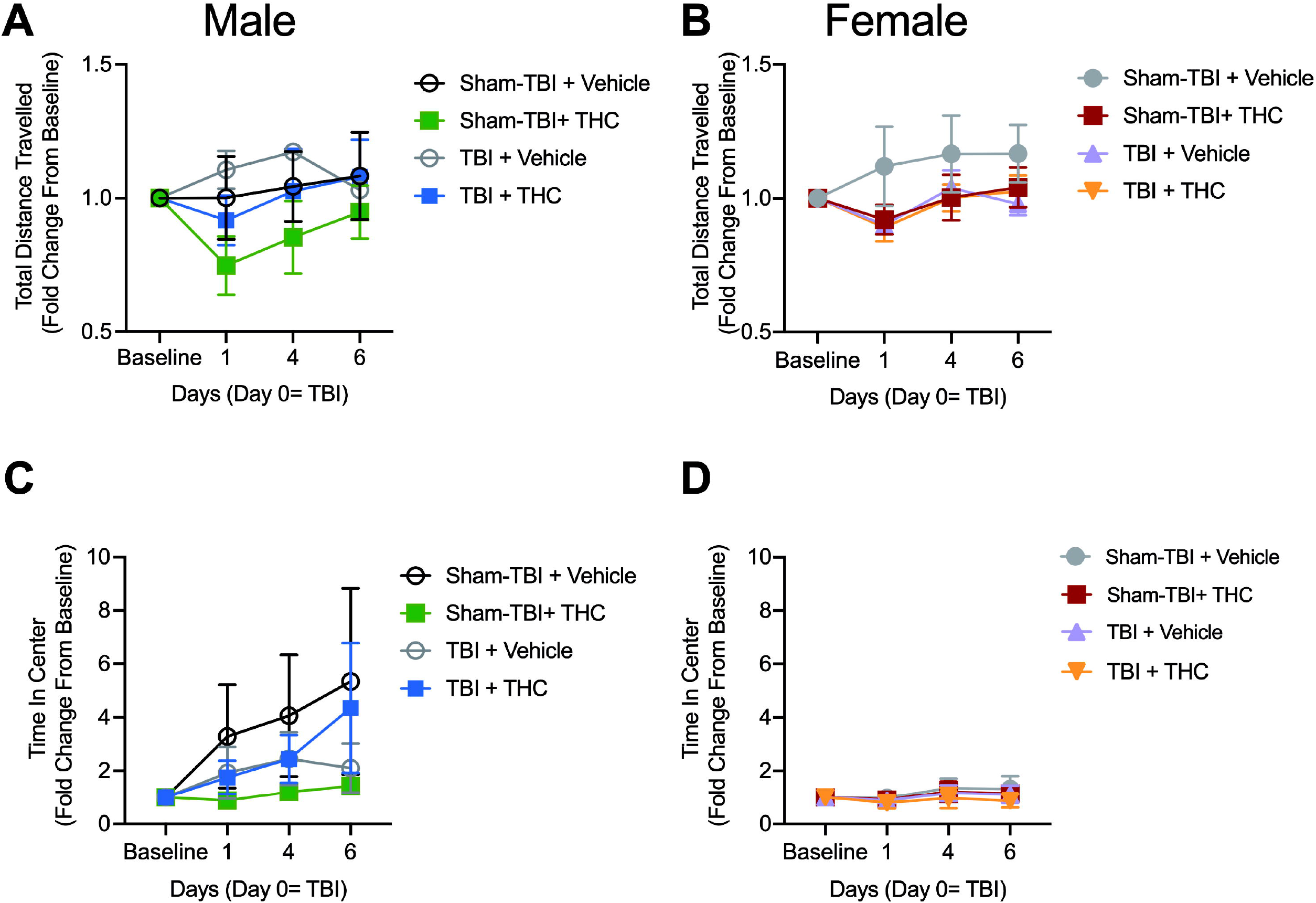
Assessment of OFT performance in rats following TBI and 1 mg·kg^-1^ THC treatment. Eighteen Sprague-Dawley rats of both sexes were administered a Sham-TBI or TBI and injected with 1 mg·kg^-1^ THC *i*.*p*. or vehicle. 1 mg·kg^-1^ THC *i*.*p*. nor TBI effected total distance travelled in males **(A)** or females **(B)** or time in the centre in male **(C)** or females **(D)**. Data presented as mean ± S.E.M. n=4-5 per group per sex. Statistical analyses were four-way RM ANOVA and described in Extended Data Table 3-1.

### Y-maze

Y-maze was used to assess changes in spatial memory on days -1 (baseline), 2 and 5 (Fig. 4; Extended Data Table 4-1). With respect to % alternations, no main effects were detected for sex, injury, drug, or interactions therein (Fig. 4A,B). A main effect of time was observed for % alternations (p = 0.048), but Tukey’s post-hoc analyses did not reveal significance within groups between timepoints (Fig. 4A,B). With respect to total arm entries, no main effects were detected for injury, drug, or interactions therein (Fig. 4C,D). Main effects of sex (p = 0.017) and time (p = 0.001) were observed for total arm entries (Fig. 4C,D); with females making more total arm entries than males. Tukey’s post-hoc analyses did not reveal significance between sexes within groups (Fig. 4C,D). Tukey’s post-hoc analyses demonstrated an increase in total arm entries for male Sham-TBI + THC animals on day 2 compared to day 5 (p = 0.013; Fig. 4C). Tukey’s post-hoc analyses did not reveal significance between timepoints within groups for females (Fig. 4D). Thus, females did enter more arms regardless of treatment and repeated exposure to the task did effect performance overall as shown by the main effect of time. Overall, neither TBI or THC changed % alternation in the y-maze, although

**Figure 4.**
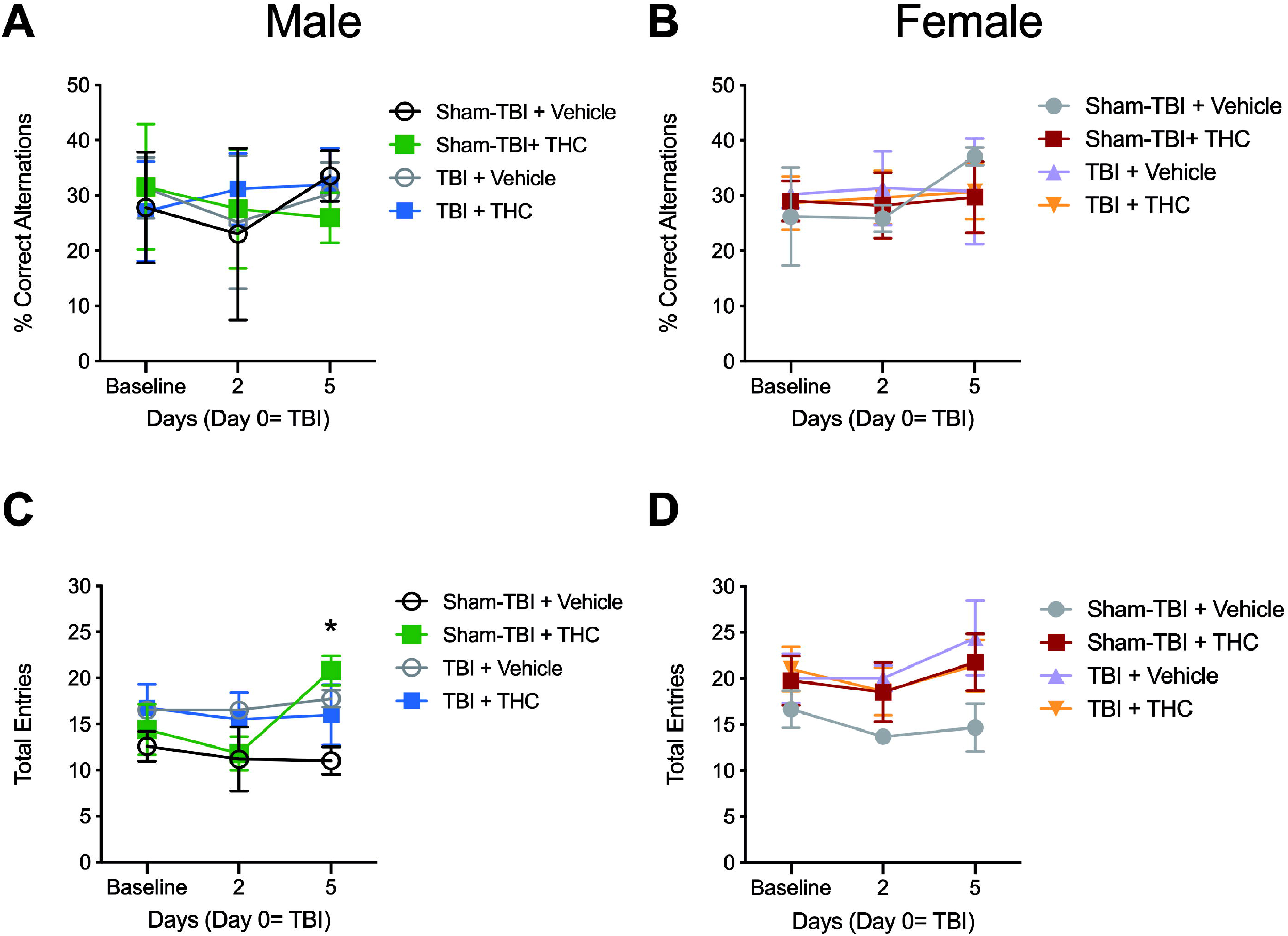
Assessment of y-maze performance in rats following TBI and 1 mg·kg^-1^ THC treatment. Eighteen Sprague-Dawley rats of both sexes were administered a Sham-TBI or TBI and injected with 1 mg·kg^-1^ THC *i*.*p*. or vehicle. 1 mg·kg^-1^ THC *i*.*p*. nor TBI effected % alternation in males **(A)** or females **(B)**. Male Sham-TBI +THC rats entered more arms on day 5 compared to day 2 compared (*p<0.05) **(C)**. 1 mg·kg^-1^ THC *i*.*p*. nor TBI effected total arm entries or females **(D)**. Data presented as mean ± S.E.M. n=4-5 per group per sex. Statistical analyses were four-way RM ANOVA and described in Extended Data Table 4-1.

### Rotarod

Rotarod was used to assess vestibulo-motor function from baseline each day of the experiment except for day 4 and 5 (Fig. 5; Extended Data Table 5-1). Latency to fall from the rotarod was recorded in seconds each day (Fig. 5A,B). No main effects were detected for injury, drug, time, or interactions therein (Fig. 5A,B). A main effect was observed for sex (p = 0.004). Tukey’s post-hoc analyses revealed a greater baseline latency to fall in the female Sham + THC animals relative to male Sham + THC animals (p = 0.043), but no other significant differences were detected (Fig. 5A,B). Given baseline variations in performance, data from the final day of testing (i.e. day 7) were expressed as fold over baseline for individual animals to determine the fold change in rotarod performance over the course of the experiment (Fig. 5C,D). A main effect of injury (p = 0.005) was observed in males, but no changes in drug or interactions were observed (Fig. 5C). Tukey’s post-hoc analysis demonstrated that latency to fall was greater for Sham-TBI + vehicle males as compared to Sham-TBI + 1 mg·kg^-1^ THC males (p = 0.049) and compared to TBI + vehicle males (p = 0.010; Fig. 5C). No changes in injury, drug, or interactions therein were detected for fold change in rotarod performance in female rats (Fig. 5D). Based on these data, 1 mg·kg^-1^ THC and TBI each hindered rotarod performance in male rats, but the combination of 1 mg·kg^-1^ THC with TBI did not improve or further limit rotarod performance. Rotarod performance was not changed by 1 mg·kg^-1^ THC or TBI in female rats.

**Figure 5.**
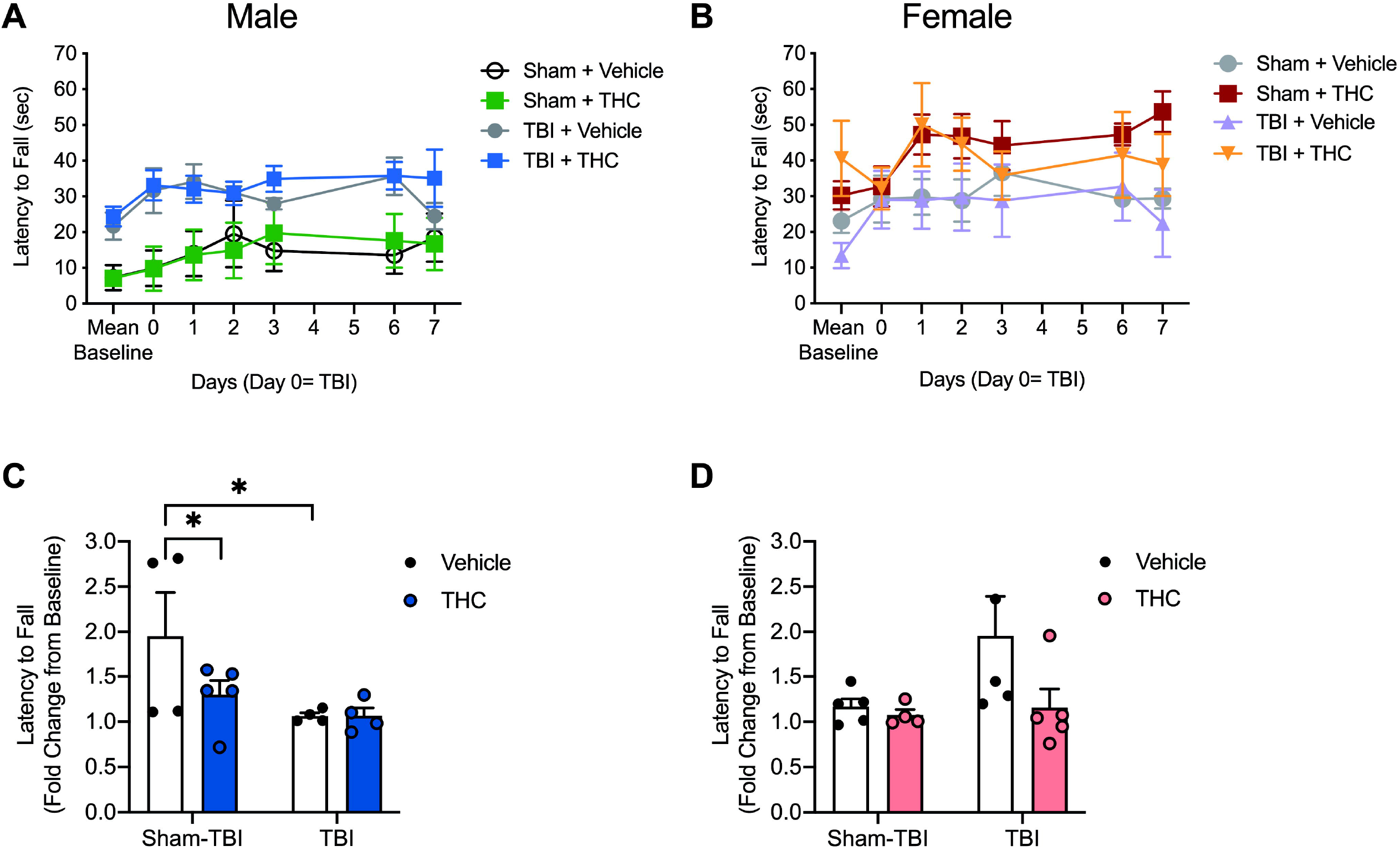
Assessment of rotarod performance in rats following TBI and 1 mg·kg^-1^ THC treatment. Eighteen Sprague-Dawley rats of both sexes were treated with a Sham-TBI or TBI and injected with 1 mg·kg^-1^ THC *i*.*p*. or vehicle. Visualization of raw data by day following treatment for males **(A)** and females **(B)**. Data were expressed as fold change from baseline over the 7-day study for males **(C)** and females **(D)**. THC decreased latency to fall versus vehicle in male Sham-TBI rats, and TBI significantly decreased latency to fall versus Sham-TBI in male rats (*p<0.05) **(C)**. There was no significant effect of 1 mg·kg^-1^ of THC or TBI on latency to fall in female rats **(D)**. Data are presented as mean ± S.E.M. fold change from baseline, n=4-5 per group (males and females). Statistical analyses were four-way RM ANOVA **(A**,**B)** or two-way ANOVA **(C**,**D)** and described in Extended Data Table 5-1.

### CB1R protein levels

Tissue samples were taken from euthanized animals on day 7 of the experiment (Fig. 1). Based on the injury location, the cortex was the area most likely effected by TBI. (Fig. 6; Extended Data Fig. 6-1, Table 6-1). Therefore, we chose to measure CB1R abundance in the cortex relative to βactin. A three-way interaction between sex, injury, and treatment was observed for CB1R levels, but no other main effects were observed (p = 0.001; Extended Data Table 6-1). In males, no main effects were detected for injury, drug, or interactions therein (Fig. 6B). In females, no main effects were detected for injury or drug; but an interaction was found between injury and drug (p = 0.003; Fig. 6C). Tukey’s post-hoc analysis revealed CB1R abundance was greater in Sham-TBI + THC females compared to both Sham-TBI + vehicle females (p = 0.030) and TBI + THC females (p = 0.008) (Fig. 6C). These data demonstrate CB1R levels increased in THC-treated females not subjected to TBI but decreased in THC-treated females subjected to TBI, whereas CB1R levels were static in males, suggesting a more dynamic regulation of CB1R abundance in female rats.

**Figure 6.**
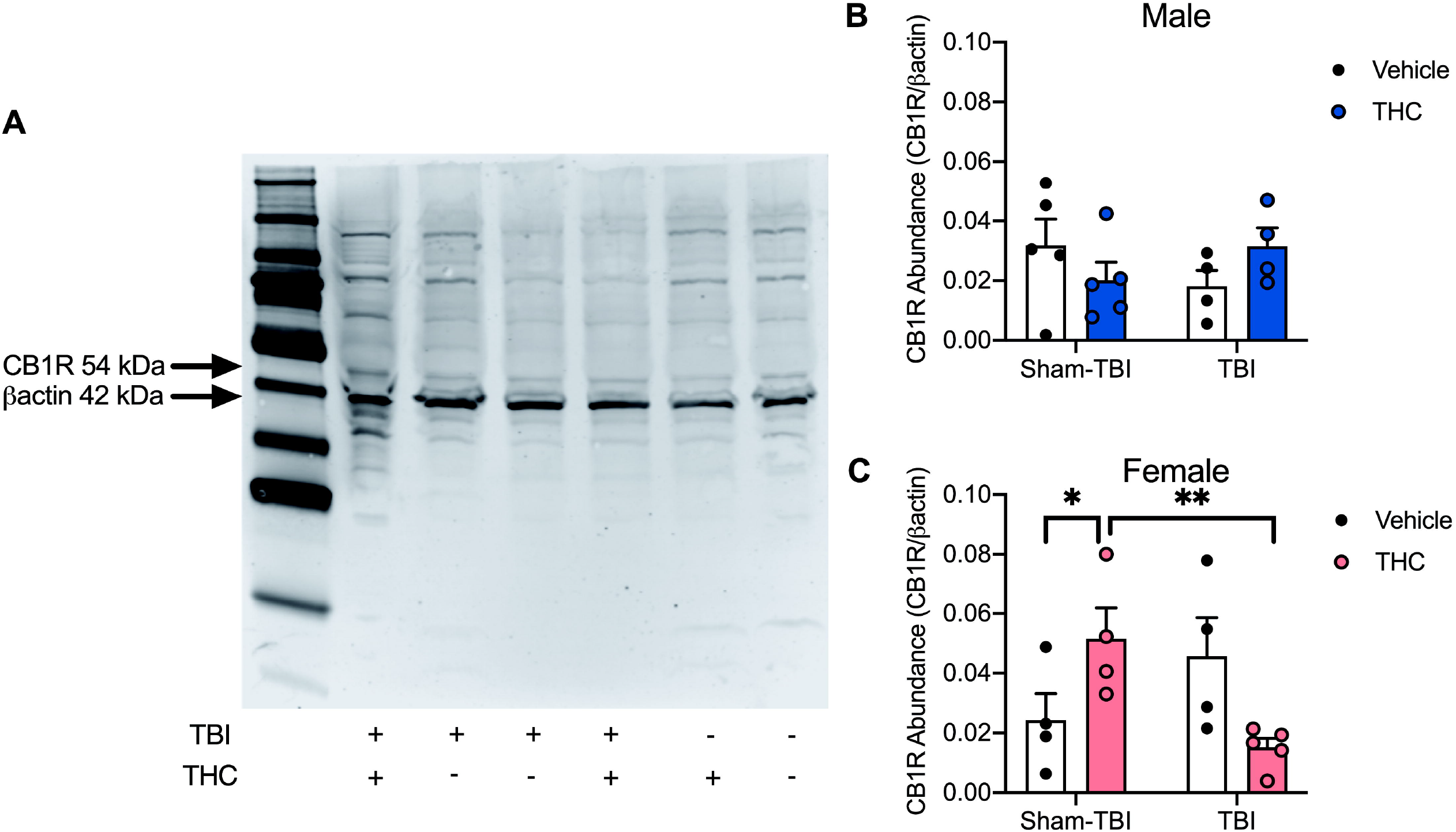
Assessment of CB1R abundance in rats following TBI and 1 mg·kg^-1^ THC treatment. Eighteen Sprague-Dawley rats of both sexes were treated with a Sham-TBI or TBI and injected with 1 mg·kg^-1^ THC *i*.*p*. or vehicle. **(A)** Representative western blot. Images were converted to 32-bit grayscale and inverted for black bands on white backgrounds. Samples were randomized. This example is rendered from a colourized full image representing male CB1R in the cortex relative to bactin. All image files used in analyses are presented in extended data figure 6-1. Densitometry data were expressed relative to bactin within sample **(B**,**C)**. Neither 1 mg·kg^-1^ of THC or TBI were associated with changes in cortical CB1R in male rats **(B)**. Treatment with 1 mg·kg^-1^ of THC and Sham-TBI was associated with higher CB1R levels compared to vehicle treatment in Sham-TBI female rats (*p<0.05) and 1 mg·kg^-1^ of THC in TBI female rats (**p<0.01) **(D)**. Data are presented as mean ± S.E.M. fold change from baseline, n=4-5 per group (males and females). Statistical analyses were three-way ANOVA and described in Extended Data Table 6-1.

### Cytokine quantification

Tissue samples were taken from euthanized animals on day 7 of the experiment (Fig. 1). Of the 27 cytokines and chemokines assessed, differences were only observed for interleukin-4 (IL-4), IL-6, CXCL5/LPS-induced chemokine (LIX), and IL-10 (Fig. 7). Data for all other cytokines and chemokines assessed can be found in extended data figure 7-1 and table 7-1. For IL-4, a three-way interaction between sex, injury, and treatment was observed, but no other main effects were present (p = 0.042; Extended Data Table 7-1). Cortical IL-4 levels were higher in TBI males compared to Sham-TBI males (p = 0.021; Fig. 7A), but no differences were observed in females (Fig. 7B). For IL-6, a main effect of injury (p = 0.043) and a three-way interaction between sex, injury, and treatment was observed, but no other main effects were present (p = 0.013; Extended Data Table 7-1). Cortical IL-6 levels were lower in Sham-TBI males treated with 1 mg·kg^-1^ THC (p = 0.010) and TBI males treated with vehicle (p = 0.043) relative to Sham-TBI males treated with vehicle (Fig. 7C). No changes in cortical IL-6 were observed in females (Fig. 7D). Finally, for LIX and IL-10, three-way interactions between sex, injury, and treatment were observed, but no other main effects were present (p = 0.037 LIX, p = 0.031 IL-10; Extended Data Table 7-1); further, post-hoc analyses did not detect any changes in cortical LIX or IL-10 levels in males or females. These data demonstrate an up- and down-regulation of IL-4 and IL-6, respectively, in our TBI model specific to male rats; these data also indicate treatment with 1 mg·kg^-1^ THC decreases IL-6 in male rats similar to TBI but the combination of THC + TBI did not reduce IL-6 in an additive manner.

**Figure 7.**
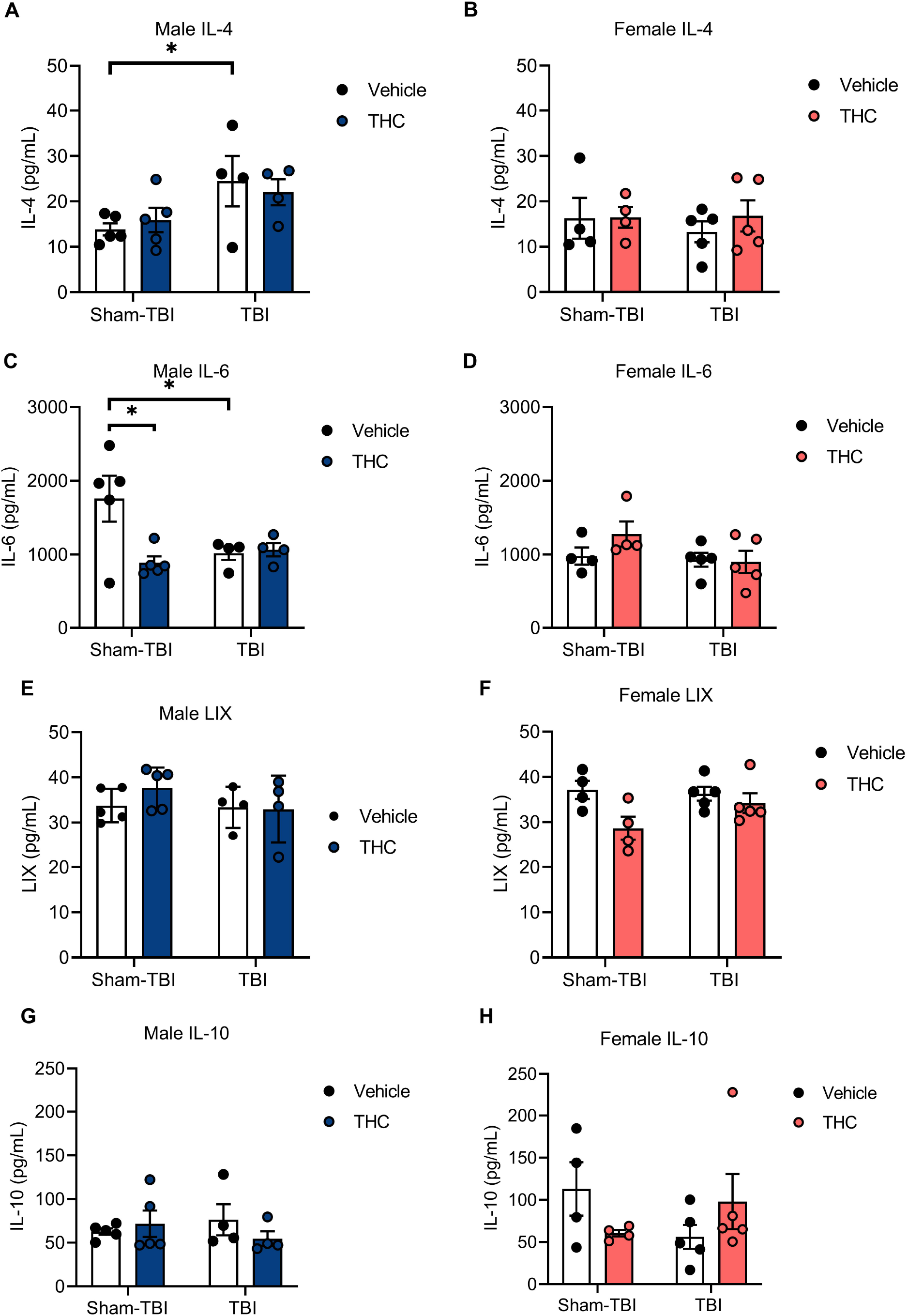
The effect of TBI and 1 mg·kg^-1^ THC on male and female IL-4 (A,B), IL-6 (C,D), LIX (E,F), and IL-10 (G,H). Eighteen Sprague-Dawley rats of both sexes were treated with a Sham-TBI or TBI and injected with 1 mg·kg^-1^ THC *i*.*p*. or vehicle. **(A)** IL-4 levels were higher in TBI + vehicle male rats compared to Sham-TBI + vehicle rats (*p<0.05). **(B)** IL-4 levels did not change in females. **(C)** IL-6 levels were lower in Sham-TBI + THC males and TBI + vehicle males (*p<0.05) compared to Sham-TBI + vehicle males. **(D)** IL-6 levels did not change in females. (**E**,**F**) LIX levels did not change in males or females. (**G**,**H**) IL-10 levels did not change in males or females. Data are presented as mean ± S.E.M. n=4-5 per group (males and females). Statistical analyses were three-way ANOVA and described in Extended Data Table 7-1.

## Discussion

Acute treatment with 1 mg·kg^-1^ THC has been shown to induce catalepsy and reduce body temperature, tail flick, and anxiety (Garai et al., 2020). In our experiments, THC was administered once, 1 h after injury; therefore, we had expected to observe tetrad effects at day 0 according to existing data (Grim et al., 2017), but did not. Anxiety-like behaviours often act as a hallmark of TBI in rodents and humans (Malkesman et al., 2013). Despite the injury magnitude being described elsewhere as a model of ‘moderate’ TBI, the injury incurred may have been too subtle to produce persistent large-scale changes of fear-based behaviour in rodents detectable in 10 min measures of OFT. To detect more subtle injury and drug induced behaviours, it is imperative to modify test parameters, for example, implementing longer recording time (>10 min), or plainly substituting more complex behaviour measures. Although y-maze has been applied in evaluating working memory for the full spectrum of TBI severity (Tucker et al., 2016; Bodnar et al., 2019), the present results indicate that it may be better suited to assess injury models that are more severe. In addition, the limited effects of THC observed here may be due to the acute and moderate dose used; future exploration of higher acute doses up to 10 mg/kg to approximate a maximal effect or repeated dosing may be used to improve this model of cannabis use after TBI.

The deficit of male rotarod performance following TBI confirms the injury model and highlights the complexities associated with comparing behavioral data between sexes. In pre-clinical research, females have been known to survive and functionally recover better in brain injury than males (Bramlett and Dietrich, 2001; Rubin and Lipton, 2019; Duncan, 2020). These sex differences have been attributed to the neuroprotective properties of progesterone and estrogen, as administration of exogenous estrogen and progesterone to males and ovariectomized females improves molecular and histological outcomes (O’Connor et al., 2005). However, outcomes seem to vary largely according to the mechanism of injury and age of rodents (Rubin and Lipton, 2019). Clinical data indicates that females are known to be at higher risk, have poorer outcomes in TBI, and are more likely to report persistent post-concussive symptoms (Farace and Alves, 2000; Dick, 2009; Bazarian et al., 2010; Alston et al., 2012). Our data demonstrated that 1 mg·kg^-1^THC had no specific beneficial effect in TBI male or female rats but was associated with decreased performance in healthy non-injured males. Moreover, IL-4 levels increased in TBI males and IL-6 levels decreased in THC and TBI males, but were unchanged in females. In contrast, cortical CB1R levels only fluctuated in females following THC treatment and TBI. The functional significance of sex-specific CB1R regulation observed in our initial study is not currently known, but suggest fundamentally different responses to TBI and THC in males and females that warrant further exploration. These preliminary data represent a single assessment of CB1R and cytokine abundance 7 days post-TBI. Additional data are required to assess dynamic changes occurring in the brain following the initial injury and recovery periods. A single dose of 1 mg·kg^-1^THC was chosen as previous reports have shown this dose to approximate an ED_50_ in rats (Zani et al., 2007; Mychasiuk et al., 2016) in acute treatment. This dose was chosen as an initial test point in these pilot studies. Future studies will evaluate higher doses of THC as well as chronic treatment paradigms. Pre-clinical research has reliably shown sex differences in cannabinoid-response; where females are more sensitive, respond faster to lower doses and exhibit higher plasma concentrations of active metabolites (Craft et al., 2012, 2017). Building from our preliminary findings, it is imperative that subsequent studies consider the importance of sex and sex hormones as contributing factors in rodent response to TBI and cannabinoids (Tseng and Craft, 2001; Craft et al., 2008, 2013; Kellert et al., 2009).

Several studies have explored the ECS and effects of cannabinoids on TBI in rodent models with several recent studies reporting elevated CB1R activity (Xing et al., 2014; Arain et al., 2015; Ponomarenko et al., 2021). In particular, one study by Bhatt et. al (2020) specifically employed THC as a pharmacological intervention in the treatment of TBI in both male and female rats. In that study, rats experienced three repeated 50 g lateral impacts to elicit diffuse repeated moderate TBIs which were either preceded by 6 doses of 1.25 mg·kg^-1^ *i*.*p*. THC (pre-injury treatment), or followed by 12 doses of 1.25 mg·kg^-1^ *i*.*p*. THC (post-injury treatment). Repeated administration of 1.25 mg·kg^-1^ *i*.*p* post-injury THC rescued working memory and reduced anxiety- and depression-like behaviour in males and females. CB1R mRNA levels were also normalized by post-injury THC treatment in the hippocampus, nucleus accumbens, and prefrontal cortex. Pre-injury treatment with THC was not associated with any benefits in this TBI model (Bhatt et al., 2020). Of note, the rats in this study were adolescents when testing occurred (post-natal day 21-55) which is problematic considering the previously highlighted roles that estrogen and progesterone can play on TBI resilience, as well as on the formation and modulation of the ECS (Craft et al., 2013). In a similar study, C57Bl/6 mice were treated with 3 mg·kg^-1^ *i*.*p*. THC after CCI and recovered rotarod performance with 2 weeks, and displayed increased levels of brain-derived neurotrophic factor and the endocannbinoid 2-arachidonoylglycerol compared to vehicle control mice that experienced CCI. (Song et al., 2021). In our study, we focused on the cortex as the primary site of injury and brain region involved in many aspects of locomotion; however, future studies should explore additional brain regions such as the hippocampus and striatum as described in Bhatt et al. and Song et al., respectively (Bhatt et al., 2020; Song et al., 2021). The comparison between our results and these previously published data suggest that THC’s effects may be greater and elicit symmetrical effect on males and females prior to sexual maturity.

From these initial data, we conclude that (i) consequences of both TBI and THC treatment are sex-dependent in rats; and (ii) in our rat model, 1 mg·kg^-1^ of THC did not overtly change TBI recovery according to the behavioral measures used. THC was selected because it is the most abundant cannabinoid found in *Cannabis sativa*, and it possesses neuroprotective and anti-inflammatory potential, but other cannabinoids such as cannabidiol also have demonstrable anti-inflammatory activity (Pertwee, 2010). These results support existing, but limited, sex differences in both TBI and ECS modulation. There is continued need for further investigation of sex-specific effects of phytocannabinoids in acute and chronic treatment paradigms relating to TBI.

## Acknowledgements

We would like to thank Anna-Maria Smolyakova for assistance with western blots, Gavin Scott for dissection instruction, and Michael Benko for help with data recording.

