## Extended Data for "Sex-specific differences in rotarod performance and type 1 cannabinoid receptor levels in a rat model of traumatic brain injury treated with Δ^9^-tetrahydrocannabinol"

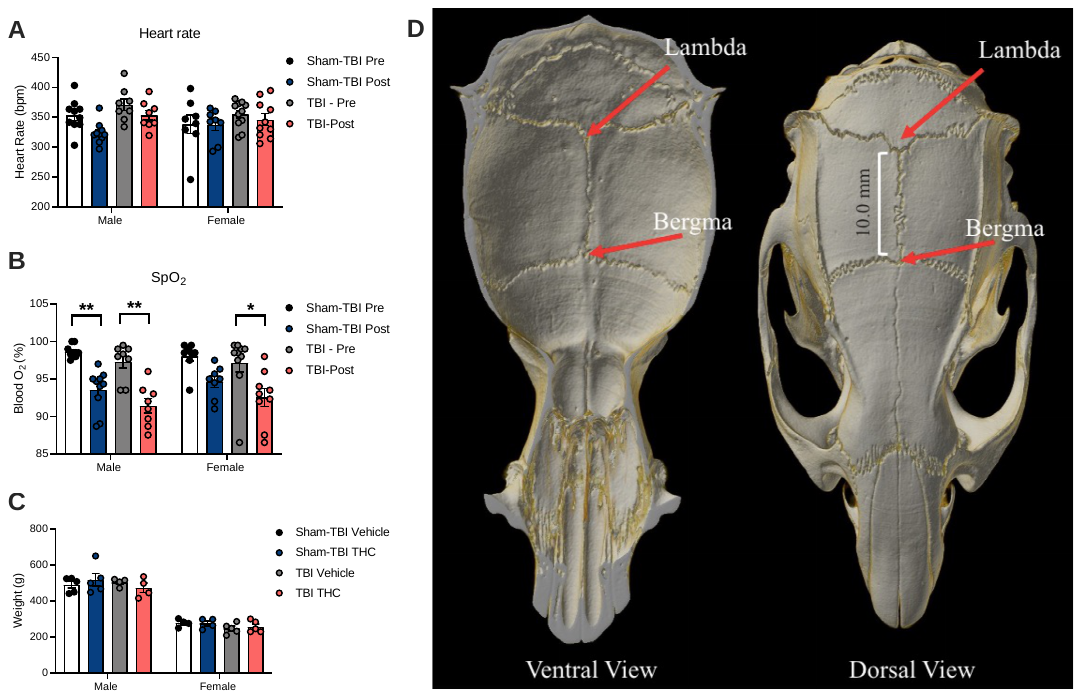

**Extended Data Figure 1-1.** Assessment of heart rate, blood oxygen saturation, weight, and skull integrity in rats following TBI.

Eighteen Sprague-Dawley rats of both sexes were administered a Sham-TBI or TBI and injected with 1 mg·kg^-1^ THC *i.p*. or vehicle (vehicle and THC treatments are combined in these data). Sham-TBI and TBI did not alter heart rate **(A)** or blood oxygen saturation **(B)** on the day of TBI. (**C**) Rat weight was not different between treatment groups 7 days post-TBI. Data presented as mean ± S.E.M. n=8-10 per group per sex. (**D**) Computed tomography scans showed no cranial fractures. *p<0.05, **p<0.01 as determined by two-way ANOVA followed by Tukey’s post-hoc test.

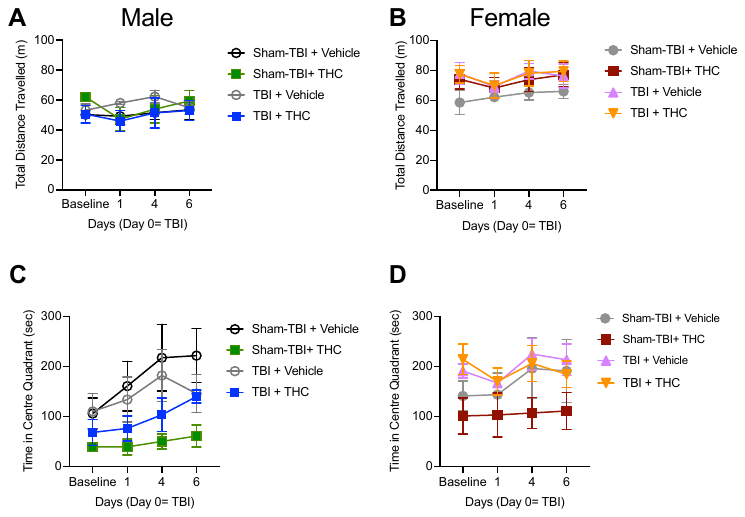

**Extended Data Figure 3-1.** Raw data for the assessment of OFT performance in rats following 1 mg·kg^-1^ THC treatment and TBI.

Eighteen Sprague-Dawley rats of both sexes were administered a Sham-TBI or TBI and injected with 1 mg·kg^-1^ THC *i.p*. or vehicle. 1 mg·kg^-1^ THC *i.p*. nor TBI effected total distance travelled in males **(A)** or females **(B)** or time in the centre in male **(C)** or females **(D)**. Data presented as mean ± S.E.M. n=4-5 per group per sex. Data were transformed as fold relative to the baseline and are presented in figure 3 with statistical analyses described in Extended Data Table 3-1.

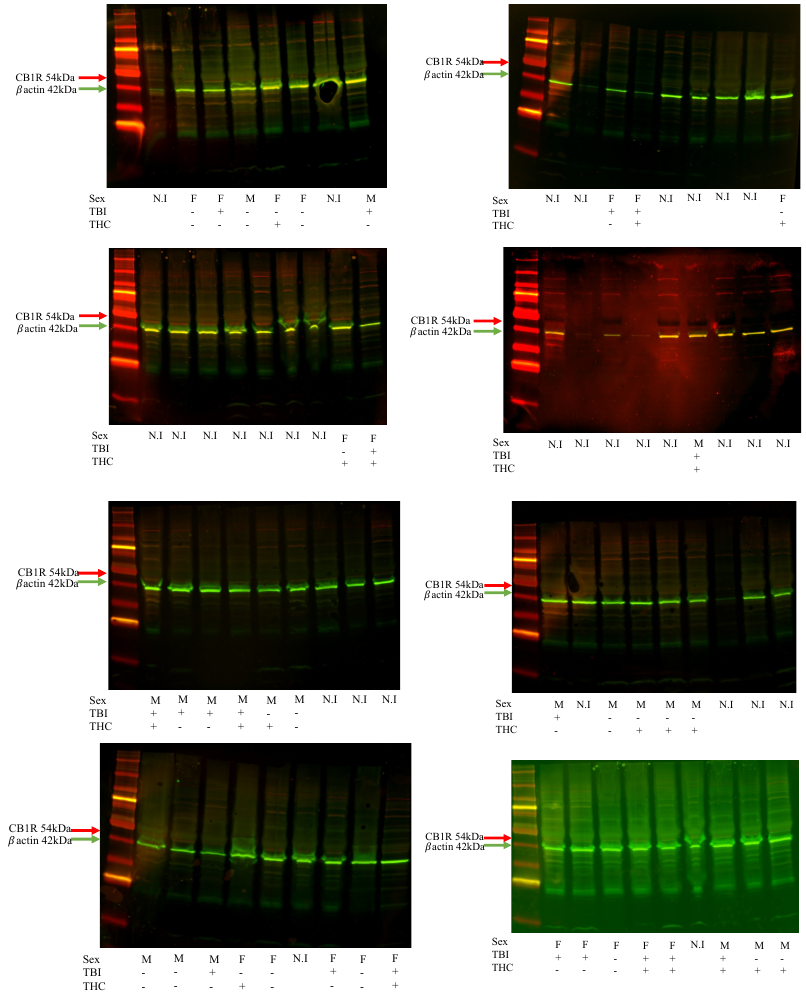

**Extended Data Figure 6-1**. All western blots used in analyses of cortex from male and female rats following 1 mg·kg^-1^ THC treatment and TBI.

n=4-5 per treatment group. N.I indicates samples that were not included in the analysis.

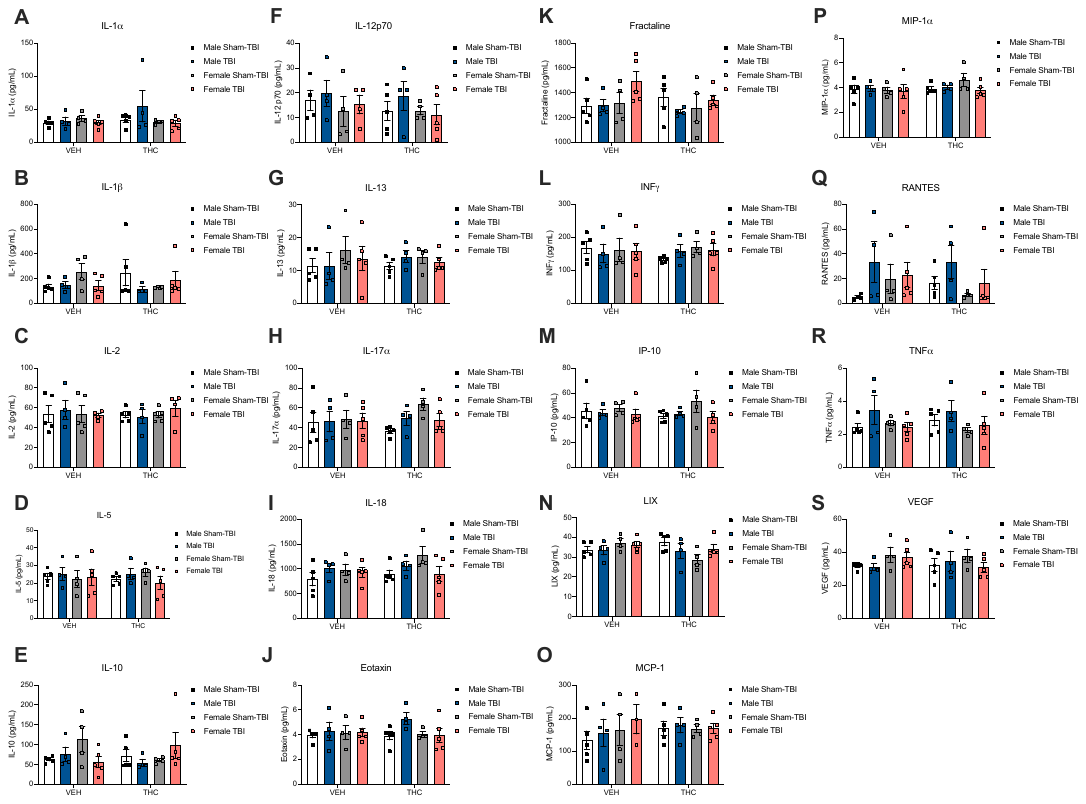

**Extended Data Figure 7-1.** The effect of 1 mg·kg^-1^ THC and TBI on male and female IL-1α **(A)**, IL-1β **(B)**, IL-2 **(C)**, IL-5 **(D)**, IL-10 **(E)**, IL-12p70 **(F)**, IL-13 **(G)**, IL-17α **(H)**, IL-18 **(I)**, eotaxin **(J)**, fractaline **(K)**, INFγ **(L)**, IP-10 **(M)**, LIX **(N)**, MCP-1 **(O)**, MIP1α **(P)**, RANTES **(Q)**, TNFα **(R)**, and VEGF **(S)**. Eighteen Sprague-Dawley rats of both sexes were treated with a Sham-TBI or TBI and injected with 1 mg·kg^-1^ THC *i.p.* or vehicle. Data are presented as mean ± S.E.M. n=4-5 per group (males and females). Statistical analyses were three-way ANOVA and described in Extended Data Table 7-1.

**Summaries of the statistical results organized by Figure. Main effects (ME) and interactions (INT) are noted throughout.**

**Extended Data Table 2-1: Tetrad response results for figure 2. Three-way 2x2x4 (injury x drug x time) repeated measures ANOVA**

|  | **ANOVA** | **F(DFn,DFd)** | **p value** |
| --- | --- | --- | --- |
| Figure 2A | ME Injury | F(1,14)=0.494 | 0.494 |
|  | ME Drug | F(1,14)=0.502 | 0.490 |
|  | ME Time | GG | 0.609 |
|  | INT | F(1,14)=0.019 | 0.893 |
| Figure 2B | ME Injury | F(1,14)=2.360 | 0.615 |
|  | ME Drug | F(1,14)=0.340 | 0.906 |
|  | ME Time | GG | 0.685 |
|  | INT | F(1,14)=0.342 | 0.499 |
| Figure 2C | ME Injury | F(1,14)=0.071 | 0.794 |
|  | ME Drug | F(1,14)=1.712 | 0.212 |
|  | ME Time | Sphericity assumed | 0.037* |
|  | INT | F(1,14)=0.115 | 0.740 |
| Figure 2D | ME Injury | F(1,14)=0.020 | 0.891 |
|  | ME Drug | F(1,14)=4.280 | 0.313 |
|  | ME Time | Sphericity assumed | 0.010* |
|  | INT | F(1,14)=0.445 | 0.516 |
| Figure 2E | ME Injury | F(1,14)=0.598 | 0.452 |
|  | ME Drug | F(1,14)=0.029 | 0.866 |
|  | ME Time | Sphericity assumed | 0.337 |
|  | INT | F(1,14)=0.003 | 0.955 |
| Figure 2F | ME Injury | F(1,14)=0.864 | 0.368 |
|  | ME Drug | F(1,14)=4.280 | 0.058 |
|  | ME Time | Sphericity assumed | 0.757 |
|  | INT | F(1,14)=0.547 | 0.472 |
| *p < 0.05; ME, main effect; INT, interaction; GG, Greenhouse-Geiser distribution in repeated measures. Assumptions of Sphericity were met according to Mauchley’s test unless indicated by GG. | | | |

**Extended Data Table 3-1: Open field test results for figure 3. Four-way 2x2x4 (sex x injury x drug x time) repeated measures ANOVA**

|  | **ANOVA** | **F(DFn,DFd)** | **p value** |
| --- | --- | --- | --- |
| Figure 3A,B | ME Sex | F(1,27)=0.233 | 0.630 |
|  | ME Injury | F(1,27)=0.021 | 0.885 |
|  | ME Drug | F(1,27)=2.417 | 0.024* |
|  | ME Time | GG | 0.099 |
|  | INT | F(1,27)=0.394 | 0.904 |
| Figure 3C,D | ME Sex | F(1,27)=4.631 | 0.040* |
|  | ME Injury | F(1,27)=2.057 | 0.163 |
|  | ME Drug | F(1,27)=6.633 | 0.016* |
|  | ME Time | GG | 0.034* |
|  | INT | F(1,27)=0.075 | 0.786 |
| *p < 0.05; ME, main effect; INT, interaction; GG, Greenhouse-Geiser distribution in repeated measures. Assumptions of Sphericity were met according to Mauchley’s test unless indicated by GG. | | | |

**Extended Data Table 4-1: Y-maze results for figure 4. Four-way 2x2x2x3 (sex x injury x drug x time) repeated measures ANOVA**

|  | **ANOVA** | **F(DFn,DFd)** | **p value** |
| --- | --- | --- | --- |
| Figure 4A,B | ME Sex | F(1,27)=0.180 | 0.675 |
|  | ME Injury | F(1,27)=0.253 | 0.619 |
|  | ME Drug | F(1,27)=0.003 | 0.959 |
|  | ME Time | Sphericity assumed | 0.048* |
|  | INT | F(1,27)=0.022 | 0.882 |
| Figure 4C,D | ME Sex | F(1,27)=6.450 | 0.017* |
|  | ME Injury | F(1,27)=3.710 | 0.065 |
|  | ME Drug | F(1,27)=1.19 | 0.285 |
|  | ME Time | Sphericity assumed | 0.001* |
|  | INT | F(1,27)=0.360 | 0.851 |
| *p < 0.05; ME, main effect; INT, interaction; GG, Greenhouse-Geiser distribution in repeated measures. Assumptions of Sphericity were met according to Mauchley’s test unless indicated by GG. | | | |

**Extended Table 5-1: Rotarod results for figure 5. Four-way 2x2x2x7 (sex x injury x drug x time) repeated measures ANOVA (figure 5A,B) or two-way 2x2 (injury x drug) ANOVA (figure 5C,D).**

|  | **ANOVA** | **F(DFn,DFd)** | **p value** |
| --- | --- | --- | --- |
| Figure 5A,B | ME Sex | F(1,27)=9.835 | 0.004* |
|  | ME Injury | F(1,27)=3.445 | 0.074 |
|  | ME Drug | F(1,27)=3.870 | 0.060 |
|  | ME Time | GG | 0.618 |
|  | INT | F(1,27)=0.02 | 0.872 |
| Figure 5C | ME Injury | F(1,14)=11.27 | 0.005* |
|  | ME Drug | F(1,14)=3.761 | 0.073 |
|  | INT | F(1,14)=3.807 | 0.071 |
| Figure 5D | ME Injury | F(1,14)=2.515 | 0.1351 |
|  | ME Drug | F(1,14)=3.244 | 0.093 |
|  | INT | F(1,14)=2.219 | 0.159 |
| *p < 0.05; ME, main effect; INT, interaction; GG, Greenhouse-Geiser distribution in repeated measures. Assumptions of Sphericity were met according to Mauchley’s test unless indicated by GG. | | | |

**Extended Data Table 6-1: Western blot results for figure 6. Three-way 2x2x2 (sex x injury x drug) ANOVA (figure 6B,C).**

|  | **ANOVA** | **F(DFn,DFd)** | **p value** |
| --- | --- | --- | --- |
| Figure 6 | ME Sex | F(1,28)=5.239 | 0.056 |
|  | ME Injury | F(1,28)=0.583 | 0.4517 |
|  | ME Drug | F(1,28)=0.813 | 0.498 |
|  | INT | F(1,28)=13.42 | 0.001* |
| Figure 6B | ME Injury | F(1,14)=0.028 | 0.870 |
|  | ME Drug | F(1,14)=0.015 | 0.905 |
|  | INT | F(1,14)=2.219 | 0.159 |
| Figure 6C | ME Injury | F(1,14)=3.054 | 0.104 |
|  | ME Drug | F(1,14)=0.808 | 0.385 |
|  | INT | F(1,14)=13.89 | 0.003* |
| *p < 0.05; ME, main effect; INT, interaction. | | | |

**Extended Data Table 7-1: Cytokine and chemokine results for figure 7. Three-way 2x2x2 (sex x injury x drug) ANOVA.**

|  | **ANOVA** | **F(DFn,DFd)** | **p value** |
| --- | --- | --- | --- |
| Figure 7A,B | ME Sex | F(1,28)=2.175 | 0.152 |
| IL-4 | ME Injury | F(1,28)=2.389 | 0.133 |
|  | ME Drug | F(1,28)=0.135 | 0.716 |
|  | INT | F(1,28)=4.545 | 0.042* |
| Figure 7C,D | ME Sex | F(1,28)=1.893 | 0.180 |
| IL-6 | ME Injury | F(1,28)=4.495 | 0.043* |
|  | ME Drug | F(1,28)=1.374 | 0.251 |
|  | INT | F(1,28)=7.046 | 0.013* |
| LIX | ME Sex | F(1,28)=0.051 | 0.823 |
|  | ME Injury | F(1,28)=0.005 | 0.944 |
|  | ME Drug | F(1,28)=1.226 | 0.278 |
|  | INT | F(1,28)=4.794 | 0.037* |
| IL-18 | ME Sex | F(1,28)=0.901 | 0.351 |
|  | ME Injury | F(1,28)=0.005 | 0.947 |
|  | ME Drug | F(1,28)=1.507 | 0.229 |
|  | INT | F(1,28)=0.636 | 0.432 |
| Eotaxin | ME Sex | F(1,28)=0.806 | 0.377 |
|  | ME Injury | F(1,28)=1.749 | 0.197 |
|  | ME Drug | F(1,28)=0.261 | 0.613 |
|  | INT | F(1,28)=0.807 | 0.377 |
| IL-1α | ME Sex | F(1,28)=1.111 | 0.301 |
|  | ME Injury | F(1,28)=0.375 | 0.545 |
|  | ME Drug | F(1,28)=0.798 | 0.379 |
|  | INT | F(1,28)=0.357 | 0.555 |
| IL-1β | ME Sex | F(1,28)=0.178 | 0.677 |
|  | ME Injury | F(1,28)=1.072 | 0.309 |
|  | ME Drug | F(1,28)=2.66 x 10^-7^ | 0.999 |
|  | INT | F(1,28)=3.694 | 0.065 |
| IL-2 | ME Sex | F(1,28)=0.023 | 0.881 |
|  | ME Injury | F(1,28)=0.161 | 0.691 |
|  | ME Drug | F(1,28)=0.011 | 0.918 |
|  | INT | F(1,28)=0.539 | 0.469 |
| IL-13 | ME Sex | F(1,28)=1.117 | 0.299 |
|  | ME Injury | F(1,28)=0.032 | 0.858 |
|  | ME Drug | F(1,28)=0.002 | 0.967 |
|  | INT | F(1,28)=0.077 | 0.784 |
| IL-10 | ME Sex | F(1,28)=1.282 | 0.267 |
|  | ME Injury | F(1,28)=0.168 | 0.685 |
|  | ME Drug | F(1,28)=0.177 | 0.677 |
|  | INT | F(1,28)=5.148 | 0.031* |
| IL-12p70 | ME Sex | F(1,28)=1.647 | 0.211 |
|  | ME Injury | F(1,28)=0.601 | 0.444 |
|  | ME Drug | F(1,28)=0.531 | 0.473 |
|  | INT | F(1,28)=0.369 | 0.549 |
| INFγ | ME Sex | F(1,28)=0.455 | 0.506 |
|  | ME Injury | F(1,28)=0.007 | 0.932 |
|  | ME Drug | F(1,28)=0.073 | 0.789 |
|  | INT | F(1,28)=0.530 | 0.473 |
| IL-5 | ME Sex | F(1,28)=0.212 | 0.648 |
|  | ME Injury | F(1,28)=0.002 | 0.963 |
|  | ME Drug | F(1,28)=0.004 | 0.949 |
|  | INT | F(1,28)=0.823 | 0.372 |
| IL-17α | ME Sex | F(1,28)=1.721 | 0.200 |
|  | ME Injury | F(1,28)=0.033 | 0.858 |
|  | ME Drug | F(1,28)=0.211 | 0.650 |
|  | INT | F(1,28)=1.495 | 0.232 |
| MCP-1 | ME Sex | F(1,28)=0.547 | 0.466 |
|  | ME Injury | F(1,28)=0.537 | 0.470 |
|  | ME Drug | F(1,28)=0.176 | 0.678 |
|  | INT | F(1,28)=0.037 | 0.850 |
| IP-10 | ME Sex | F(1,28)=0.469 | 0.499 |
|  | ME Injury | F(1,28)=1.579 | 0.220 |
|  | ME Drug | F(1,28)=0.016 | 0.901 |
|  | INT | F(1,28)=0.690 | 0.413 |
| VEGF | ME Sex | F(1,28)=2.006 | 0.167 |
|  | ME Injury | F(1,28)=0.424 | 0.521 |
|  | ME Drug | F(1,28)=0.083 | 0.776 |
|  | INT | F(1,28)=0.645 | 0.429 |
| TNFα | ME Sex | F(1,28)=3.113 | 0.087 |
|  | ME Injury | F(1,28)=1.713 | 0.201 |
|  | ME Drug | F(1,28)=3.67 x 10^-4^ | 0.985 |
|  | INT | F(1,28)=0.692 | 0.413 |
| RANTES | ME Sex | F(1,28)=0.611 | 0.441 |
|  | ME Injury | F(1,28)=3.880 | 0.059 |
|  | ME Drug | F(1,28)=0.086 | 0.771 |
|  | INT | F(1,28)=0.388 | 0.539 |
| Fractaline | ME Sex | F(1,28)=1.319 | 0.261 |
|  | ME Injury | F(1,28)=0.397 | 0.534 |
|  | ME Drug | F(1,28)=0.699 | 0.410 |
|  | INT | F(1,28)=0.001 | 0.979 |
| MIP-1α | ME Sex | F(1,28)=0.019 | 0.891 |
|  | ME Injury | F(1,28)=0.527 | 0.474 |
|  | ME Drug | F(1,28)=1.172 | 0.288 |
|  | INT | F(1,28)=0.625 | 0.436 |
| *p < 0.05; ME, main effect; INT, interaction. | | | |
